# Effect of AM114 on apoptosis and proinflammatory cytokines in IL-1-beta treated human THP-1 derived macrophages

**DOI:** 10.1101/2023.05.08.539939

**Authors:** Chitra Selvarajan, Vasanth Konda Mohan, Nalini Ganesan, MK Kalaivani

## Abstract

**Background:** Chalcones and their derivatives are precursors of flavonoids and isoflavonoids abundantly present in edible plants. It displays a wide range of pharmacological activities, such as anticancer, antiinflammatory, antibacterial and antioxidant. AM-114 (3,5-*Bis*-[benzylidene-4-boronic acid]-1-methylpiperidin-4-one), a boronic-chalcone derivative exhibits potent anticancer activity through inhibition of the proteasome on human colon cancer cell line. However, the cytotoxic and anti-inflammatory effects of AM114 on THP-1 derived macrophages remain to be studied.

**Aim of the study:** Inflammation is a complex reaction managed by a variety of immune cells such as monocytes and macrophages. In response to inflammatory stimuli, the macrophages secrete increased proinflammatory cytokines, chemokines such as interleukin-6 (IL-6) and interleukin-8(IL-8). The proteasome complex is essential for several cellular processes including protein degradation, cellular differentiation and antigen presentation. In this study, human monocyte cell line THP-1 stimulated with interleukin-1-beta (IL-1-beta) was used as a model to investigate the *in vitro* effects of AM114, a proteasome inhibitor (PI), on apoptosis and release of proinflammatory cytokines.

**Materials and methods:** The effect of AM114, with or without IL-1-beta stimulated THP-1derived macrophages, was assessed by comparing with aspirin. The cell viability was determined by MTT assay. The qualitative measurement of apoptosis was determined by acridine orange/ ethidium bromide (AO/EtBr), Hoechst 33342 staining and rhodamine 123 assays. The quantitative measurement of apoptosis was carried out to determine the caspase-3 activity by spectrofluorimetric method and DNA fragmentation by agarose gel electrophoresis. The release of proinflammatory cytokines and chemokines such as IL-6 and IL-8 was also measured by enzyme linked immunosorbent assay (ELISA) method. The ANOVA was used to compare the different groups.

**Results:** In the present study the IC_50_ concentration of AM114 and Aspirin on THP-1 cells were determined as 3.6 micromolar and 3900 micromolar respectively. The THP-1 derived macrophages treated with IL-1-beta showed insufficient apoptosis and increased release of proinflammatory cytokines. However the treatment of AM114, on stimulated THP-1 derived macrophages showed pronounced apoptotic features. The caspase-3 activity was increased 1.9 fold in AM114 with IL-1-beta treated THP-1 derived macrophages as compared to the unstimulated THP-1 derived macrophages pretreated with AM114. An increase of 3.2 fold in caspase-3 activity was observed in stimulated THP-1 derived macrophages pretreated with AM114 in comparison with stimulated THP-1 derived macrophages pretreated with aspirin. A significant increase of 5.0 and 3.0 fold DNA damage were observed in stimulated THP-1 derived macrophages treated with AM114 and aspirin as compared to unstimulated cells. Moreover AM114 treated cells showed significant decrease (p value less than 0.001) in the release of IL-6 and IL-8 on stimulated THP-1 derived macrophages.

**Conclusion:** The results on the induction of apoptosis and suppression of proinflammatory cytokines suggest that AM114 is more effective than aspirin in stimulated THP-1 derived macrophages.

## 1. Introduction

Inflammation is a vital part of the complex biological response of body tissues to various harmful stimuli and plays an important role in the advancement of a wide array of diseases such as bacterial sepsis, cardiovascular diseases and rheumatoid arthritis (RA) (Chen et al., 2021). One of the most important soluble mediators of inflammation is interleukin-1-beta (IL-1-beta) a potent proinflammatory cytokine produced by blood monocytes and mediates wide range of reactions involved in acute phase responses. Monocytes/macrophages play a central role in both, contributing to the final consequence of chronic inflammation which is represented by the loss of tissue function due to fibrosis. The activated macrophages are the key inflammatory cells, plays an important role in the inflammatory diseases by secreting the proinflammatory cytokines such as IL-1-beta, Tumor necrosis factor (TNF)-_α_ and interleukin-6 (IL-6). Therefore targeting the macrophage mediated inflammatory responses is necessary to develop a novel therapeutic approach against inflammatory conditions.

The ubiquitin-proteasome pathway has a central role in the selective degradation of intracellular proteins. Among the key proteins modulated by the proteasome are those involved in the control of inflammatory processes. Consequently proteasome inhibition is a potential treatment option for cancer and inflammatory conditions. Thus far, proof of principle has been obtained from studies in numerous animal models for a variety of human diseases including inflammatory conditions such as rheumatoid arthritis, asthma, multiple sclerosis, psoriasis and cancer.

Our earlier review article on proteasome inhibitor also demonstrated the ubiquitin proteasome system and efficacy of proteasome inhibitors in diseases (Chitra et al., 2012). Aspirin or acetylsalicylic acid (ASA) is a nonsteriodal anti-inflammatory drug effective in treating fever, pain and inflammation in the body. It inhibits the proteasome function in RA and induces abnormalities in mitochondria, which plays a critical role in the stimulation of apoptotic signals and anti-inflammatory responses (Dikshit et al., 2006).

Chalcones and their derivatives are precursors of flavonoids and isoflavonoids abundantly present in edible plants. It display a wide range of pharmacological activities, such as antimalarial, anticancer, antiprotozoal (antileishmanial and antitrypanosomal), antiinflammatory, antibacterial, antifilarial, antifungal, antimicrobial, larvicidal, anticonvulsant and antioxidant. AM-114 (3,5-*Bis*-[benzylidene-4-boronic acid]-1-methylpiperidin-4-one), a boronic-chalcone derivative exhibits potent anticancer activity through inhibition of the proteasome on human HCT 116 colon cancer cell line (Michalkova et al., 2023). However, so far no study has reported the cytotoxic effect of AM114 on THP-1 derived macrophages.

In the present study THP-1 monocyte cell lines were used to mimic the primary monocyte derived macrophages. The IL-1-beta proinflammatory cytokine was used as a stimulant to mimic the inflammatory condition in RA. The main objective of the study was to explore the effect of AM114 on apoptosis and the release of proinflammatory cytokines in IL-1-beta stimulated THP-1 derived macrophages.

## 2. Materials and Method

### 2.1 Materials

Roswell Park Memorial Institute Medium (RPMI) 1640, 10% fetal calf serum (FCS), 25 mM HEPES, 2 mM L-glutamine, MTT [3-(4,5-dimethylthiazol-2-yl)-2,5-diphenyl tetrazolium bromide] were purchased from Invitrogen, EU. Penicillin, streptomycin, kanamycin, fungizone were from Himedia (Mumbai, India). AM114 was purchased from Tocoris biosciences, MO, USA. Agarose, Tris Triton X-100 EDTA (TTE), Sodium chloride (NaCl), isopropanol, ethanol, Dimethyl sulfoxide (DMSO), Rhodamine 123 and Hoechst 33342 were sourced from Sigma Aldrich (MO, USA). Human THP-1 monocytes cells were procured from national centre for cellular sciences (NCCS) (Pune, India). Human recombinant IL-1-beta (rh IL-1-beta) was purchased from R&D systems (MN). EnzChek® Caspase-3 Assay Kit was purchased from Molecular probes invitrogen (EU). Human IL-6 and IL-8 ELISA kits were procured from Krishgen biosystems (Mumbai.India).Other reagents and chemicals were of the highest purity grade.

### 2.2 Cell culture

Human THP-1 monocytes were maintained in suspension at a density of 2 ×10^5^ / mL to 1×10^6^ / mL in RPMI 1640 media containing 10 % fetal calf serum. To differentiate THP-1 monocytes to macrophages the cell suspensions 1×10^5^ / mL were plated in RPMI 1640 media with 10% FCS, 5% human serum containing L-glutamine 2 mmol/L, HEPES 25mM, L-glutamine 2 mM, penicillin 100 IU/mL, streptomycin 100 mg/mL, kanamycin 20µg/mL, fungizone 20 µg/mL at 37 ° C with 5% CO_2_.

### 2.3 MTT assay

To determine the toxicity of AM114 and Aspirin on THP-1 cells, the cell viability was measured by 3-(4,5-dimethylthiazol-2-yl)-2,5-diphenyl tetrazolium bromide (MTT) assay method as described by McKenzie (2019). Determining the IC_50_ values for THP-1 derived macrophage viability in culture was a prerequisite that was carried out in a 96 well plate. Plates were seeded with 10^3^ cells/ mL of cells and treated with 0-5.0 µM of AM114 or 0-10,000 µM of aspirin in triplicates. Viability of cells after 24 h was determined using MTT (10 µL) reagent and incubated further for 4 h. Dimethyl sulphoxide (100 µL) solubilization was done and absorbance measured with a microtitre plate reader (FLUO STAR OPTIMA BMG LABTECH) at 650 nm. Percent viability was calculated as (OD of drug treated sample/control OD) × 100.

### 2.4 *In vitro* cell stimulation treatment

Adhered cells cultured in RPMI, 10% FCS with 5% human serum were pretreated with AM114 and Aspirin for 45 min and then stimulated with human IL-1-beta (10 ng / mL) for 24 h. The induction of apoptosis in AM114 and aspirin treated macrophages with or without stimulation for 24 h were observed using (i) AO/EtBr fluorescence staining (ii) Hoechst 33342 assay and (iii) Rhodamine 123 assay.

### 2.5 Qualitative measurement of apoptosis

#### 2.5.1 Acridine orange/ Ethidium bromide (AO/EtBr) staining

The effects of AM114 and aspirin on apoptosis in THP-1 cells were determined by AO/EtBr dual staining according to the protocol described previously by Brousseau et al., 1999. After 24 hrs of drug treatment, cell media was removed and washed twice with PBS. 1 µL of dye mixture (100 mg/mL acridine orange (AO) and 100 mg/mL ethidium bromide (EtBr), in distilled water) was added per well and incubated for 1min at room temperature. After staining the cells were then observed under an inverted phase contrast fluorescent microscope at ×200 magnifications (Nikon Eclipse, USA) with an excitation filter at 480 nm.

#### 2.5.2 Hoechst 33342 staining

Hoechst 33342 was employed to label both intact and apoptotic nuclei. The changes in the nucleus were detected by Hoechst 33342 staining assay as described previously by Thangam et al., 2014. Cells were seeded onto 24 well plates at a density of 1×10^5^ cells / well. Following treatment, the cells were washed in ice-cold phosphate buffered saline (PBS; pH 7.4), fixed with 4% (w/v) paraformaldehyde and incubated with (10 mg/mL) Hoechst 33342 for 3 minutes at room temperature. Condensed and fragmented nuclei were evaluated by intercalation of the fluorescent probe into nuclear DNA.

The stained nuclei were visualised in blue channel fluorescence with an inverted phase contrast fluorescent microscope at ×200 magnifications.

#### 2.5.3 Measurement of mitochondrial membrane potential by Rhodamine 123 efflux assay

The changes in mitochondrial membrane potential were measured by Rhodamine 123 staining assay as described by Thangam et al., 2014. For the Rhodamine 123 assay, after 24 hr drug treatment culture medium was discarded and the cells were washed twice with 1 mL PBS. The cells were fixed in 4% formaldehyde and incubated for 20 min at room temperature. After incubation the plates were washed with methanol followed by PBS. The Rhodamine 123 stain (1mg/10mL) was added and incubated at room temperature for 20 minutes. After incubation cells are rinsed with methanol to remove the excess stain and washed with PBS. The mitochondrial depolarization patterns of drug treated and control cells under inverted phase contrast fluorescent microscope at ×200 magnifications were observed with an excitation and emission wavelengths of 505nm and 534nm, respectively.

### 2.6 Quantitative measurement of apoptosis

#### 2.6.1 Caspase -3 activity assay

The caspase-3 activity was measured using the EnzChek Caspase-3 Assay Kit according to manufacturer’s protocol. Briefly, the cells 1×10^5^ cells / tube were lysed for 10 min on ice in 50 µL of the lysis buffer provided by the manufacturer. After centrifugation (5000 g at 4 °C for 5 minutes), 50 µL lysates were incubated with caspase-3 substrate at 37 °C for 30 minutes. Fluorescence was measured in a fluorescence microplate reader (HYBRID MULTIMODE) using excitation/emission wavelength of 496/520 nm.

#### 2.6.2 Measurement of DNA damage by DNA fragmentation method

The DNA damage by apoptosis was analysed by DNA fragmentation by agarose gel electrophoresis based on the method of Rahbar Saadat et al., 2015. The THP-1 cells genomic DNA was extracted by adding 0.5 mL of lysis buffer (TTE solution), incubated for 1.5 h in 37 ° C then centrifuged at 10,000 rpm/4° C/10 min. The supernatant was discarded and 0.5 mL of TTE solution was added followed by the addition of 100 µL NaCl and vortexed. Then 0.7 mL of isopropanol and 0.1 mL 5 M NaCl was added. Tubes were incubated overnight at -20° C then centrifuged again at 10,000 rpm/RT/15 min afterwards the DNA pellets were rinsed by adding 500 µL of ice-cold 70% ethanol and centrifuged at 10,000 rpm for 10 minutes at 4 ° C. The DNA pellets were dissolved by adding 20-50 µL of Tris EDTA (TE) solution. Ten micro liter of DNA sample was mixed with 3 µL loading buffer/ lane were run on 1% agarose gel to determine the DNA damage levels. After electrophoresis, the gel was stopped and the DNA was photographed under GelDoc System (UVP).

### 2.7 Measurement of IL-6 and IL-8

The cells were plated into six well plates and treated with AM114 and aspirin for 45 mins. After pretreatment, the cells were treated with IL-1-beta for 24 h. The IL-6 and IL-8 in culture supernatants were detected by ELISA by following the manufacturer’s instructions.

### 2.8 Statistical analysis

The experiments were carried out in triplicates and the data were expressed as mean ± standard error mean (SEM). The IC50 values were calculated using Graph pad Prism 6 software. One-way analysis of varience (ANOVA) was calculated to compare the experimental groups by Graph Pad Instat 3 software.

## 3. Results

### 3.1 Cytotoxic effects of AM114 and aspirin on THP-1 cells

The effect of AM114 and Aspirin on viability of THP-1 derived macrophages was studied. Cells were incubated for 24 h with different doses of AM114 concentrations, ranging from 0 to 5 µM and aspirin concentrations ranging from 0 to 10,000 _μ_M, and cell viability was determined by MTT assay. As seen in Fig. 1a and 1b a dose dependent decrease in cell viability was observed. The mean IC_50_ of these dose–response studies for AM114 and aspirin was found to be 3.6 µM and 3900 _μ_M respectively.

**Fig. 1.**
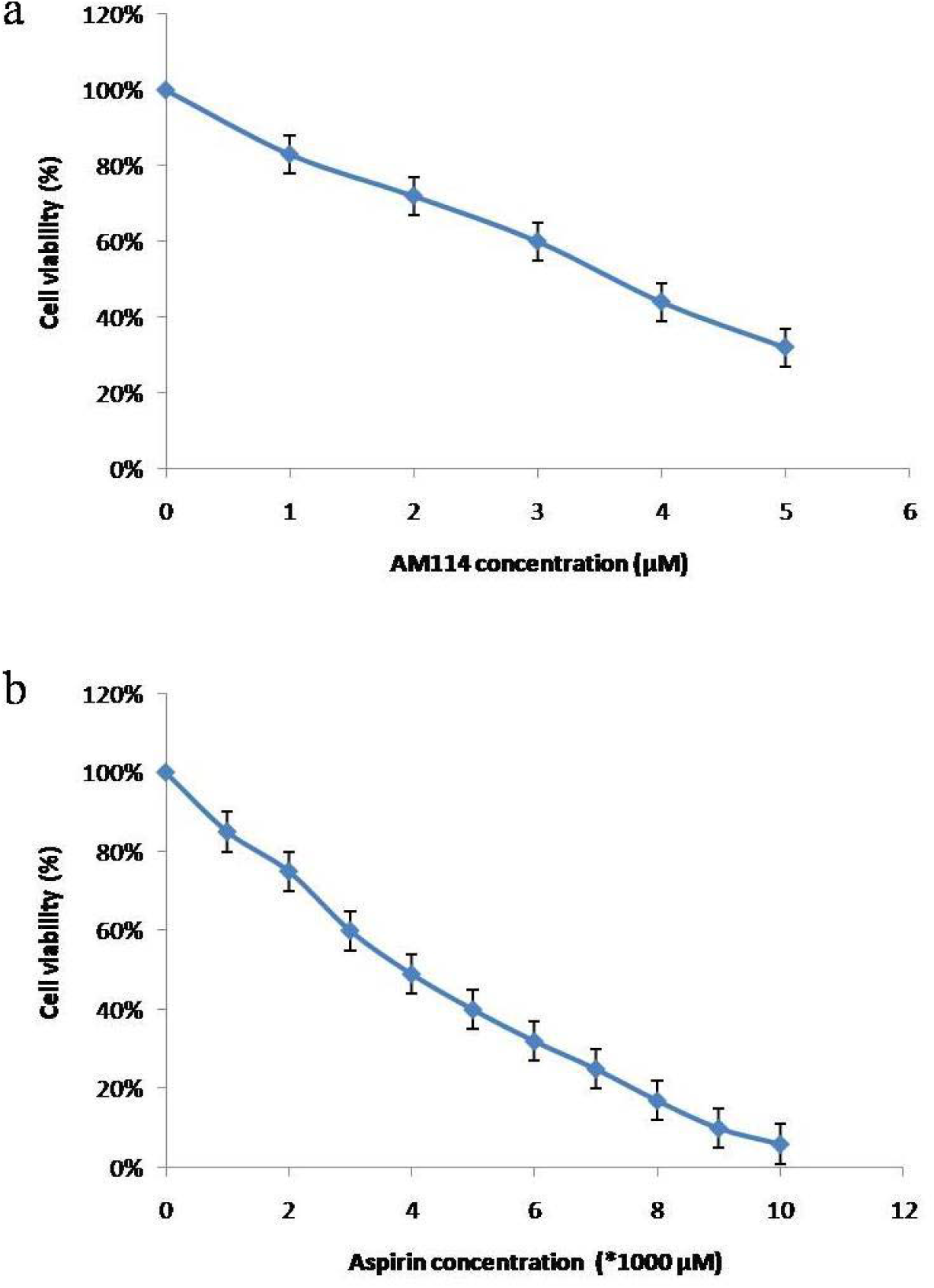
MTT assay on THP-1 derived macrophages. Cytotoxic effect of (a) AM114 (IC_50_ :3.6 _μ_M) and (b) Aspirin (IC_50_ : 3900 _μ_M) on THP-1 cells. Data are presented as mean± SEM of triplicate samples.

### 3.2 Induction of apoptosis by AM114 on THP-1 derived macrophages

Fig. 2 showed the fluorescence staining of AO/EtBr assay. The fluorescence images revealed that the cells stained green represent viable cells, whereas yellow staining represented early apoptotic cells, and reddish or orange staining represents late apoptotic cells. In the control, uniformly green live cells with normal and large nucleus were observed (Fig. 2a). The macrophages stimulated with IL 1-beta alone did not show any apoptotic changes (Fig. 2d). Where as in AM114 and aspirin treated cells yellow, orange and red staining was observed to a lesser extent (Fig. 2b and 2c) when compared to the drug treated IL-1-beta stimulated macrophages (Fig. 2e and 2f). The increased apoptosis were found to occur in AM114 with stimulation than the Aspirin with stimulation (Fig. 2e and Fig. 2f). The characterization of the cell death induced by AM114 and aspirin was further examined with the help of fluorescent DNA binding agent, Hoechst 33342. These results confirm that AM114 and aspirin significantly induced the apoptosis in stimulated THP-1 derived macrophages.

**Fig. 2.**
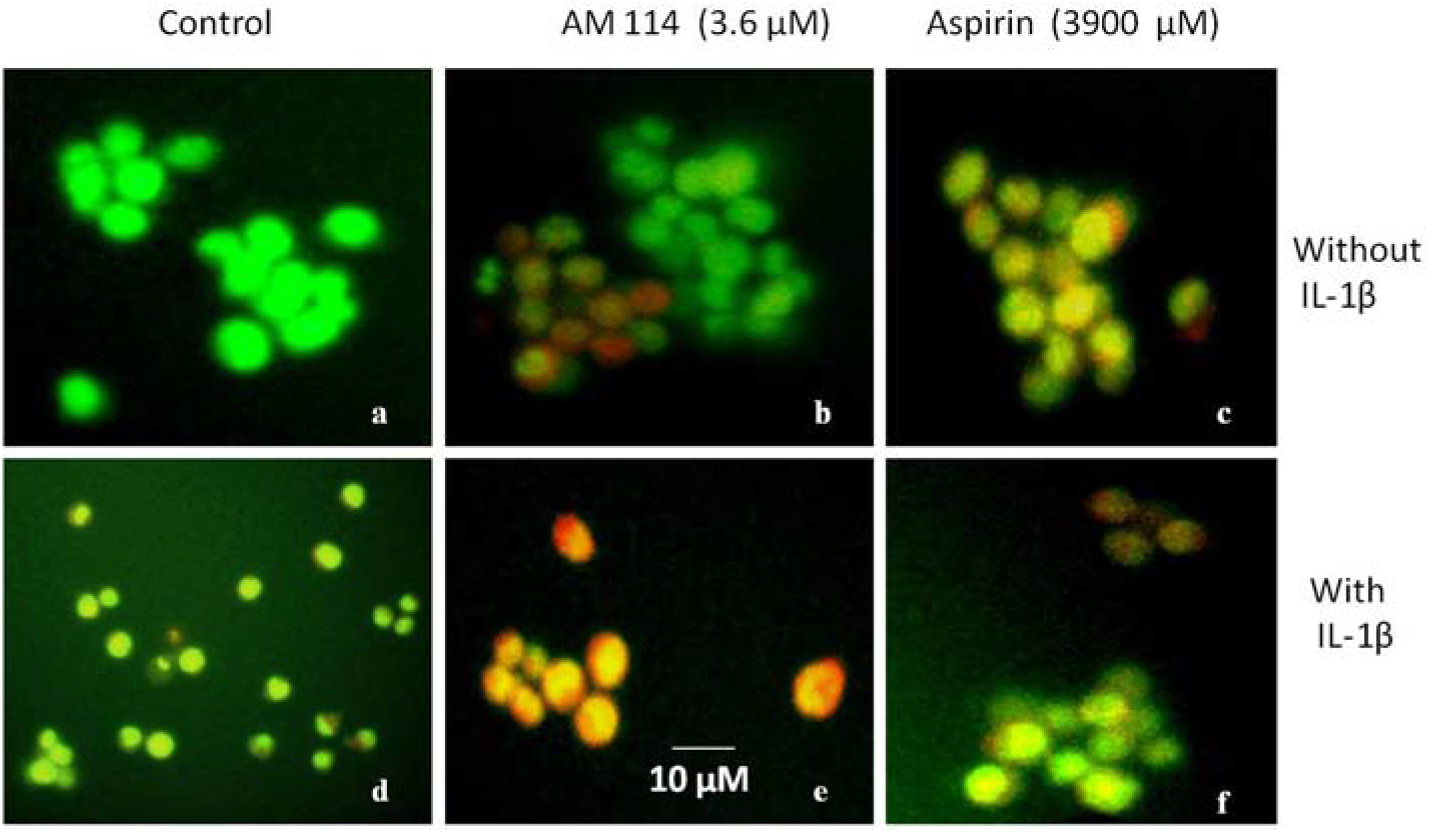
AO/EtBr assay on THP-1 derived macrophages. Fluorescence microscopy images of THP-1 derived macrophages exposed to AM114 (IC50: 3.6 _μ_M) and aspirin (IC50: 3900 _μ_M) for 45 min followed by stimulation with and without IL-1-beta (10 ng/mL) for 24 h. The panel of images shows AO/EtBr staining for induction of apoptosis. Top panel: without IL-1-beta (a) control; (b) AM114; (c) aspirin. Bottom panel: with IL-1-beta (d) control; (e) AM114; (f) aspirin. Scale: 10 _μ_m.

Staining the nucleus with blue fluorescence Hoechst33342 showed the induction of apoptosis upon treatment of cells with AM114 and aspirin with or without stimulation for 24h (Fig. 3). The unstimulated THP-1 derived macrophages showed normal cell morphology (Fig. 3a). In contrast clear induction of apoptosis was seen in AM114 and aspirin treated cultured macrophages (Fig. 3b and Fig. 3c). The cells treated with IL-1-beta alone did not induce apoptosis of macrophage cells (Fig. 3d). The increased apoptotic characteristics such as membrane blebbing, fragmented nucleus, and condensed chromatin and apoptotic bodies were found to occur in AM114 with stimulated cells when compared to aspirin with stimulation (Fig. 3e and Fig. 3f).

**Fig. 3.**
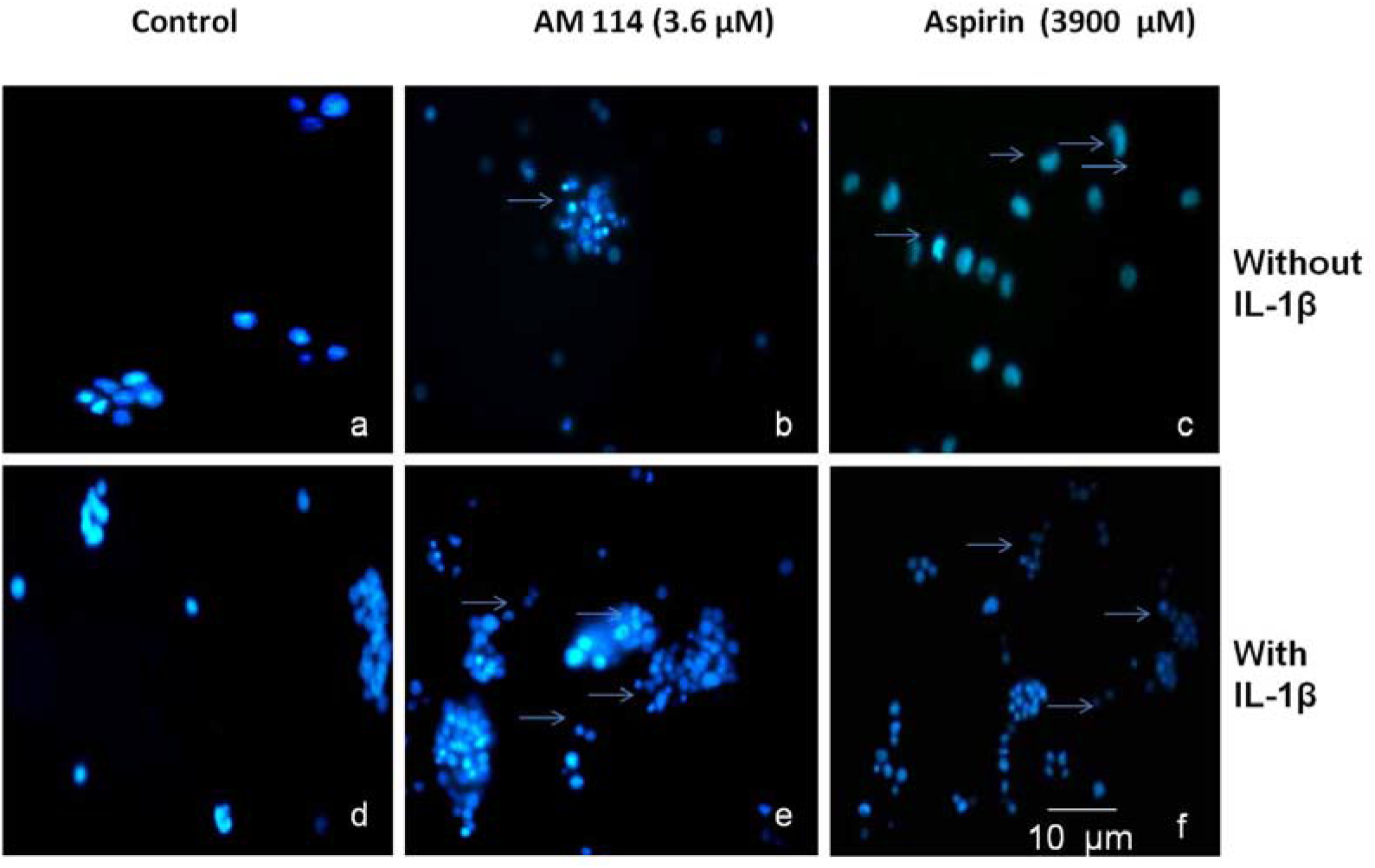
Hoechst 33342 assay on THP-1 derived macrophages. Fluorescence microscopy images of THP-1 derived macrophages exposed to AM114 (IC50: 3.6 _μ_M) and aspirin (IC50: 3900 _μ_M) for 45 min followed by stimulation with and without IL-1-beta (10 ng/mL) for 24 h. The panel of images shows Hoechst 33342 staining indicates nuclear fragmentation. Top panel: without IL-1-beta (a) control; (b) AM114; (c) aspirin. Bottom panel: with IL-1-beta (d) control; (e) AM114; (f) aspirin. Scale: 10 _μ_m.

Fig. 4 showed the fluorescence images of rhodamine 123 assay. In THP-1 derived macrophages, rhodamine dye aggregates in mitochondria as a function of membrane potential, resulting in red fluorescence with brightness proportional to the membrane potential (Fig. 4a). In contrast less mitochondrial membrane changes with decreased red fluorescence in few cells was seen in AM114 and aspirin treated cultured macrophages (Fig. 4b and 4c). The cells treated with IL-1-beta alone showed few bright red fluorescent cells indicate the normal function of the mitochondrial membrane (Fig. 4d). In contrast IL-1-beta stimulation showed increased mitochondrial membrane changes with reduced red fluorescence in large number of cells treated with the two drugs (Fig. 4e and 4f).

**Fig. 4.**
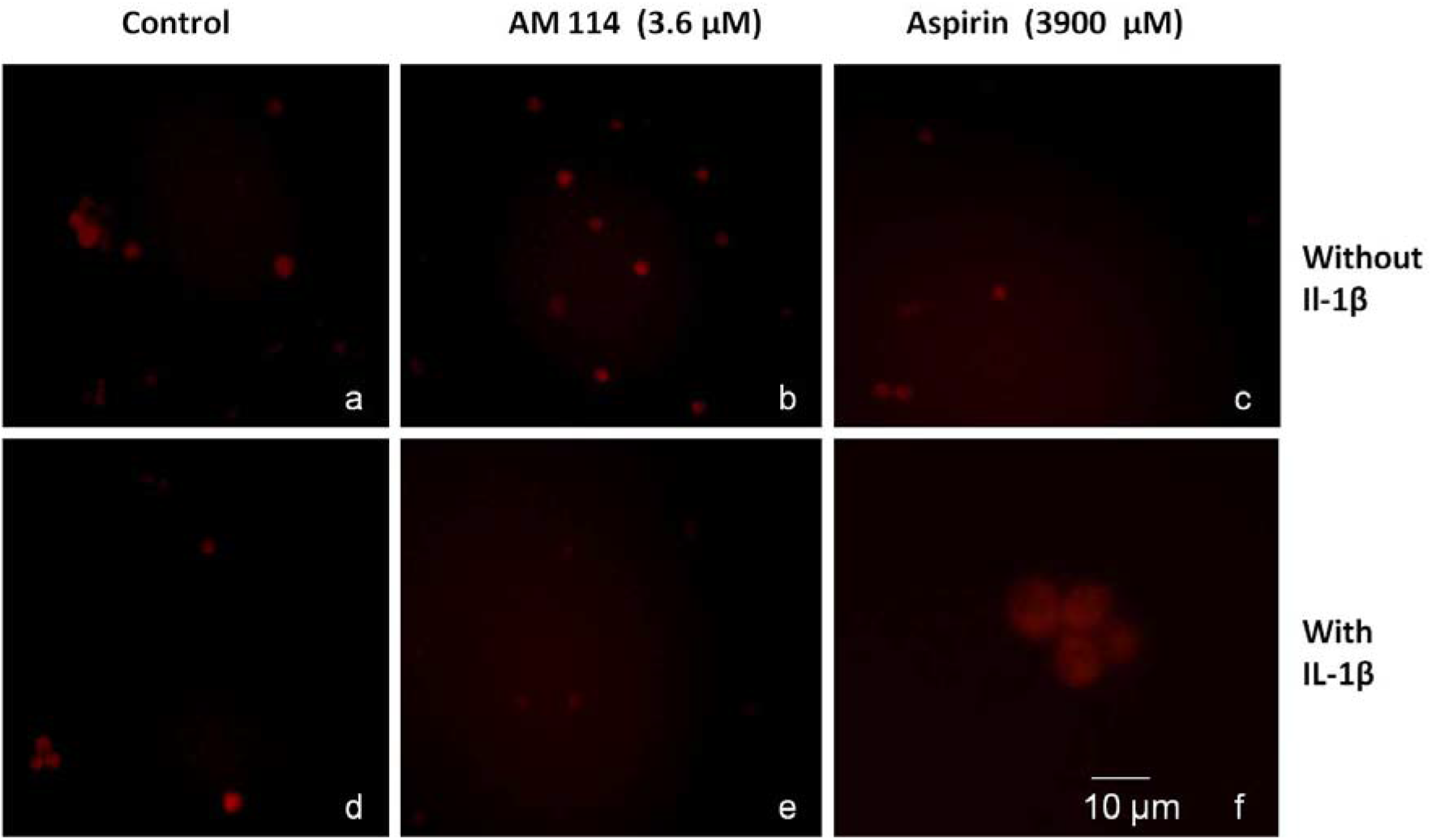
Rhodamine 123 assay on THP-1 derived macrophages. Fluorescence microscopy images of THP-1 derived macrophages exposed to AM114 (IC50: 3.6 _μ_M) and aspirin (IC50: 3900 _μ_M) for 45 min followed by stimulation with and without IL-1-beta (10 ng/mL) for 24 h. The panel of images shows Rhodamine 123 assay staining indicates changes in mitochondrial membrane potential. Top panel: without IL-1-beta (a) control; (b) AM114; (c) aspirin. Bottom panel: with IL-1-beta (d) control; (e) AM114; (f) aspirin. Scale: 10 _μ_m.

The results of caspase-3 activity in AM114 and aspirin treated macrophages with or without stimulation are shown in Fig. 5. The caspase-3 activity in IL-1-beta treated macrophages decreased by 0.2 fold than unstimulated THP-1 derived macrophages. In the presence of IL-1-beta the AM114 showed 1.9 fold increased caspase-3 activities than the AM114 without stimulation. The aspirin with IL-1-beta treated macrophages showed 0.6 fold elevated caspase-3 activity than the aspirin without stimulation. An increase of 3.2 fold caspase-3 activity was observed in AM114 stimulated cells in comparison with aspirin stimulated cells.

**Fig. 5.**
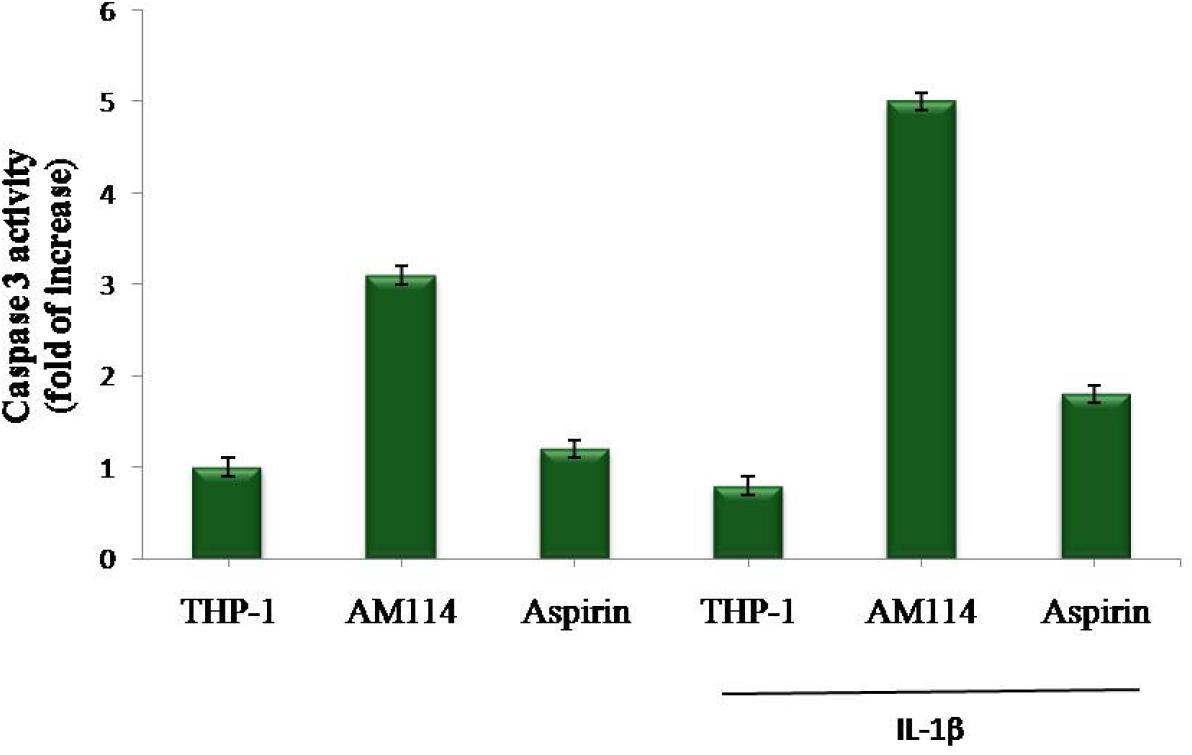
Caspase-3 activity assay on THP-1 derived macrophages. Caspase-3 activity in AM114 (IC50: 3.6 _μ_M) and aspirin (IC50: 3900 _μ_M) treated THP-1 derived macrophages with and without IL-1-beta (10 ng/mL) for 24 h. The results are mean ± standard error mean of triplicate cultures.

The extent of DNA damage by apoptosis on THP-1 derived macrophages was shown in Fig. 6. Fig. 6(a) represents the DNA damage by DNA fragmentation assay on THP-1 derived macrophages. Fig. 6(b) showed the histogram of DNA damage percent represented in fold increase. In THP-1 derived macrophages, the DNA damage increased 1.4 fold in IL-1-beta stimulated macrophages (Lane 2) when compared to unstimulated macrophages (Lane 1). Treatment with AM114 and aspirin unstimulated macrophages, showed 2.8 and 1.8 fold increased DNA damage (Lane 3 & Lane 4) when compared with THP-1 derived macrophages (Lane 1). A significant increase of 5.0 and 3.0 fold DNA damage (Lane 5 & Lane 6) were observed in stimulated macrophages treated with AM114 and aspirin as compared to THP-1 derived macrophages (Lane 1).

**Fig. 6.**
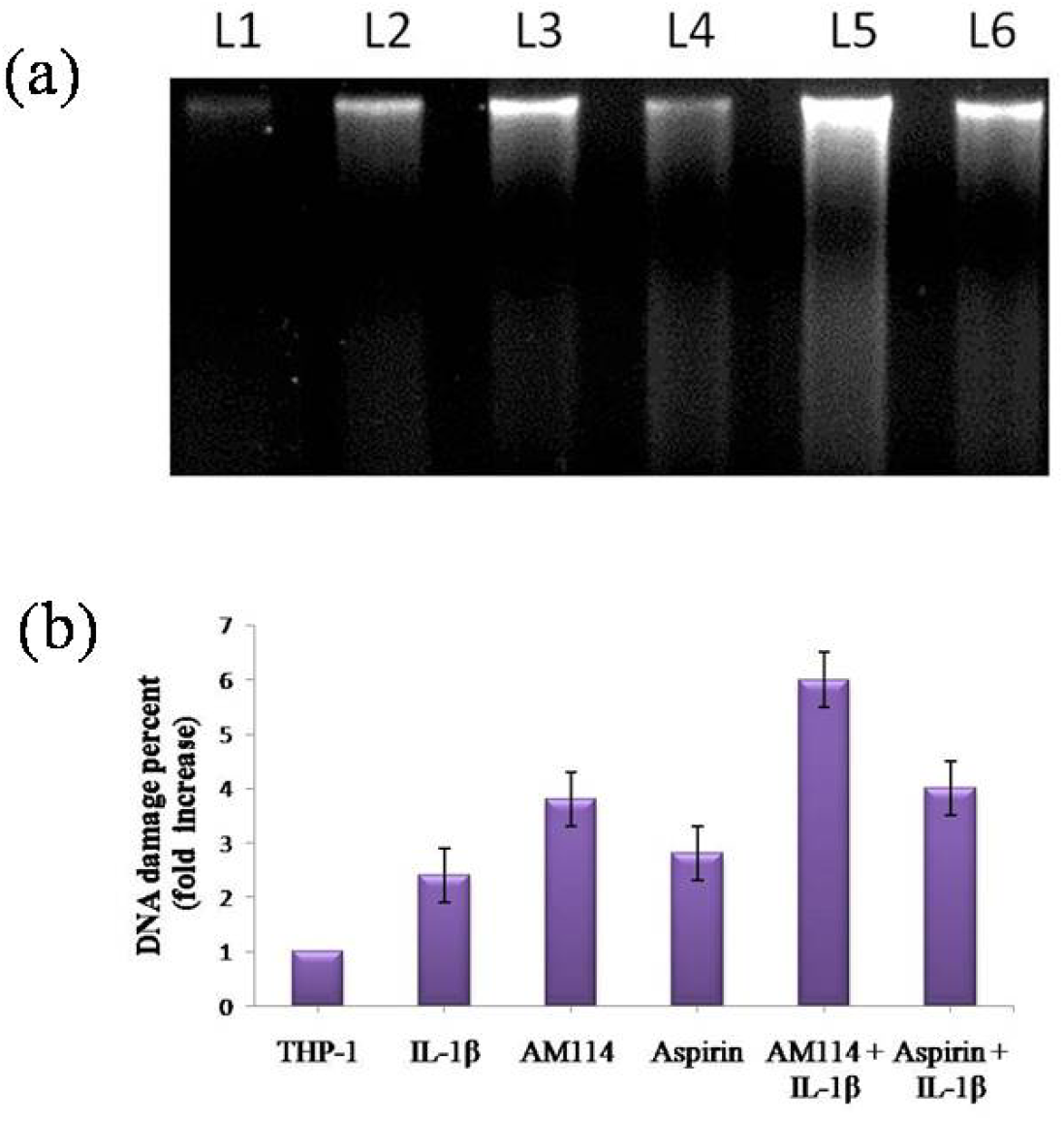
DNA damage by DNA fragmentation on THP-1 derived macrophages. (a) The DNA damage on 1% agarose gel electrophoresis. Lane 1 -THP-1 derived macrophages; Lane 2 - IL-1-beta stimulated THP-1 derived macrophages; Lane 3 - AM114 treated THP-1 derived macrophages; Lane 4 - Aspirin treated THP-1 derived macrophages; Lane 5 - AM114 with IL-1-beta stimulated THP-1 derived macrophages; Lane 6 - Aspirin with IL-1-beta stimulated THP-1 derived macrophages. (b) The histogram of DNA damage percent on THP-1 derived macrophages represented in fold increase.

In order to assess the effect of AM114 and aspirin on the release of proinflammatory cytokine and chemokine related to the inflammatory reaction, the levels of IL-6 and IL-8 were measured in THP-1 derived macrophages stimulated with or without IL-1-beta. Fig. 7 shows the measurement of IL-6 cytokine on THP-1 derived macrophages. The levels of IL-6 were significantly increased (p<0.001) in the IL-1-beta stimulated THP-1macrophages when compared to unstimulated THP-1 derived macrophages. The pretreatment of AM114, aspirin in unstimulated macrophages showed no significant changes as compared to the unstimulated macrophages of THP-1. However, the pretreatment with AM114, aspirin in stimulated macrophages showed highly significant suppression in the levels of (p<0.001) IL-6 cytokine as compared to the stimulated macrophages of THP-1 without pretreatment. Similar results but of lesser magnitude was observed in aspirin treated stimulated cells.

**Fig. 7.**
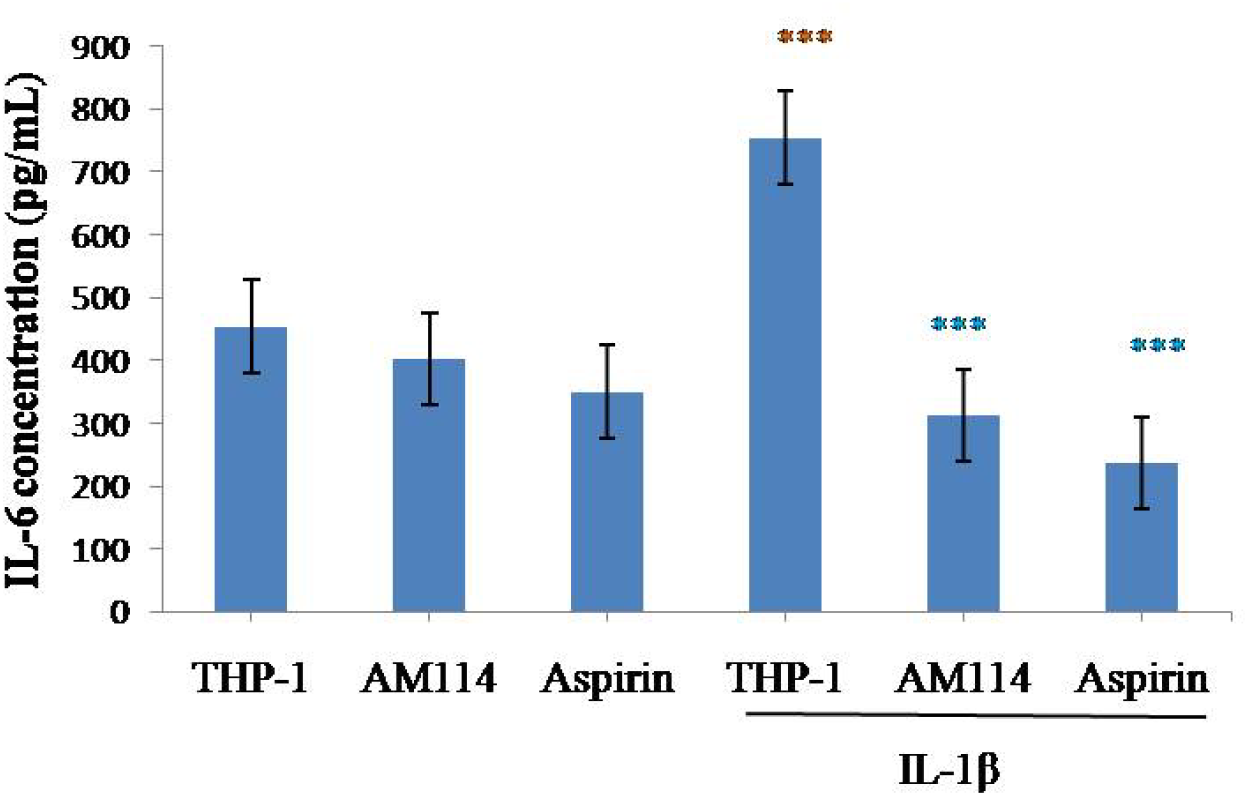
Effect of AM114 at IC_50_ concentration on the production of IL-6 cytokine in the THP-1 derived macrophages. Macrophages were pretreated with AM114 and aspirin for 45 min and further incubated with and without IL-1-beta (10 ng/mL) for 24 h. IL-6 production was measured in culture supernatants by ELISA method. Data represents mean ± SEM of triplicates. ***p<0.001 - IL-1-beta stimulated THP-1 derived macrophages Vs Unstimulated THP-1 derived macrophages; ***p<0.001 - Stimulated macrophages pretreated with AM114, aspirin Vs Stimulated THP-1 derived macrophages.

Fig. 8 shows the measurement of IL-8 chemokine on THP-1 derived macrophages. The levels of IL-8 were significantly increased (p<0.001) in the IL-1-beta stimulated THP-1 derived macrophages when compared to unstimulated THP-1 derived macrophages. The pretreatment of AM114, aspirin in unstimulated macrophages showed no significant changes as compared to the unstimulated macrophages of THP-1. However, the pretreatment with AM114, aspirin in stimulated macrophages showed highly significant suppression in the levels of (p<0.001) IL-8 cytokine as compared to the stimulated macrophages of THP-1 without pretreatment. Similar results but of lesser magnitude was observed in aspirin treated stimulated cells.

**Fig. 8.**
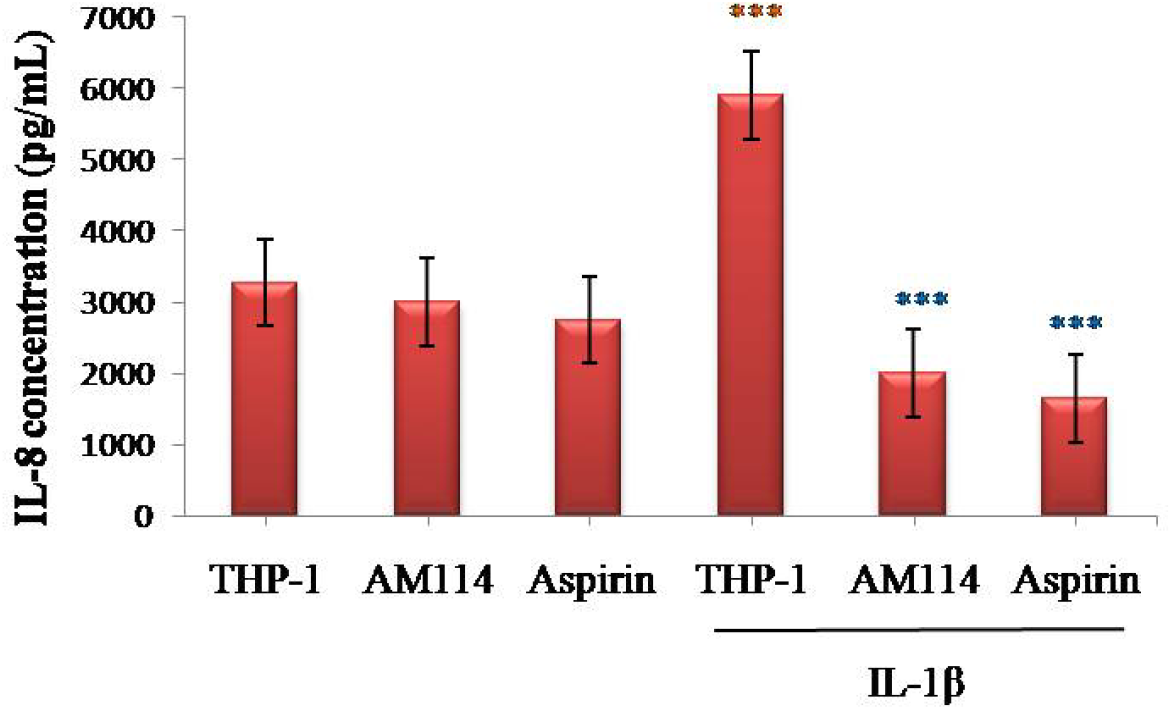
Effect of AM114 at IC50 concentration on the production of IL-8 chemokine in THP-1 derived macrophages. Macrophages were incubated with AM114 and aspirin for 45 min and further incubated with and without IL-1-beta (10 ng/mL) for 24 h. IL-8 production was measured in culture supernatants by ELISA. ***p<0.001 - IL-1-beta stimulated THP-1 derived macrophages Vs Unstimulated THP-1 derived macrophages; ***p<0.001 - Stimulated macrophages pretreated with AM114 and aspirin Vs Stimulated THP-1 derived macrophages.

## 4. Discussion

Our earlier studies on macrophages demonstrated the comparison of differentiation to macrophages in isolated human peripheral blood and THP-1 cells (Chitra et al., 2014). The cell growth and size was optimized and demonstrated in macrophages derived from human peripheral blood and THP-1 cells. The proteasome has a central role in the regulation of macrophage function by inhibiting the proteasome with several naturally occurring inhibitors (Qureshi et al., 2012). The efficient therapies against TNF-_α_ and IL-1-beta have identified macrophages as a crucial target for therapeutic intervention (Ponzoni et al., 2018).

In the present study, AM114 and Aspirin exerted cytotoxic effect in a dose dependent manner on THP-1 cells. The IC_50_ concentration of AM114 and Aspirin was found to be 3.6 µM and 3900 µM in THP-1 cells respectively. It was reported earlier that AM114 and aspirin exerted cytotoxic effect on macrophages derived from peripheral blood of RA patients at IC_50_ concentration of 0.88 µM and 1120 µM respectively (Chitra et al., 2016). The studies by Castano et al., 1999 and Dikshit et al., 2006 explored the cytotoxicity of aspirin against human adenocarcinoma HT-29 cell line and mouse neuro 2a neuroblastoma cells with IC_50_ concentration of 5 mM and 2.5 mM. Nibret et al., 2021 tested the *in vitro* cytotoxicity of AM114 against human HCT 116 colon cancer cell line with IC_50_ of 1.5 µM.

The insufficient apoptosis of inflammatory cells might contribute to the pathogenesis in inflammatory diseases such as RA. Our earlier study on RA macrophages has explored the efficacy of AM114 on the apoptosis induction in stimulated RA macrophages (Chitra et al., 2016). Aspirin and other NSAIDs induce apoptosis in several cell types such as human colorectal tumour cell line, fibroblast cell line, B cell chronic lymphocytic leukemia cells and myeloid leukemia cell lines (Piazza et al., 1999; Bellosillo et al., 1998; Castaño et al., 1999; Shiff et al., 1995; Lu et al., 1995; Schwenger et al., 1997; Klampfer et al., 1999).

It has been reported that the cells undergoing apoptosis exhibit several morphological and cellular changes including cytoplasmic blebbing, nuclear shrinkage, chromatin condensation, irregularity in shape, DNA fragmentation and cleavage of key cellular proteins (Kari et al., 2022). The studies by Kerr, Winterford, & Harmon (1994) have shown similar morphological changes in the cancer cells that undergo apoptosis. Aspirin induces apoptosis through the inhibition of proteasome function. It has been documented that aspirin induced mitochondrial membrane depolarization in neuro 2a cells (Dikshit et al.,2006). It has been reported earlier that bortezomib induced the induction of apoptosis and inhibition of T cell activation in RA patients (Ghannam et al., 2020). The proteasome inhibitor lactacystin, induced apoptosis on human monoblastic U937 cells (Imajoh-Ohmi et al., 1995).

Caspase-3 played key roles in the post-mitochondrial apoptotic pathway including the activation of several caspases which are the hallmarks of apoptosis (Boatright et al., 2003; Zhao et al., 2013). The caspase-3 when activated has many cellular targets that produce the morphologic features of apoptosis (McIlwain et al., 2013). It functions as major effector by cleavage of numerous cell death substrates, to cellular dysfunction and destruction (Riedl and Shi, 2004).

The studies by Dikshit et al.(2006) demonstrated the induction of apoptosis by proteasome inhibitor aspirin in THP-1 cells, neuro 2a cells Jurkat, and HL-60, though the release of cytochrome c from mitochondria and activation of caspases. Taburee et al. (2011) have demonstrated the involvement of caspase-3 pathway in cytokine induced apoptosis in leukemic cell line. A biochemical hallmark of apoptosis was the cleavage of chromatin into small fragments including oligonucleosomes, which were described as DNA ladders in the electrophoresed gel (Archana et al., 2013). The aspirin inhibits the proteasome function and causes severe mitochondrial abnormalities, playing critical role in the induction of apoptotic signal and anti-inflammatory responses (Dikshit et al.,2006).

Our study results also confirm the similar apoptotic features in AM114 and Aspirin treated stimulated THP-1 derived macrophages. In addition, our results also showed significant increased levels of caspase-3 and DNA fragmentation in AM114 and aspirin with IL-1-beta - treated cells. This may be due to impairment in proteasome function and mitochondrial abnormalities with subsequent release of caspase-3. AM114 pretreatment increased the DNA fragmentation as evidences by agarose gel electrophoresis indicating the effectiveness of the drug which proves that AM114 potentially exerts the cell destruction and enhanced apoptosis in inflammation.

In our study, the pretreatment of AM114 showed suppression in the release of IL-6, IL-8 in IL-1-beta stimulated macrophages indicates that AM114 has significant anticytokine potential that would be most beneficial in inflammatory disease conditions such as RA. In this respect, our data are consistent with the earlier reports by Van der Heijden et al. (2009) who explored the effect of bortezomib, in suppressing the production of proinflammatory cytokines in activated T cells of RA. Similarly the proteasome inhibitor TPCK suppressed the release of proinflammatory cytokine in cultured monocytes/macrophages (Rovin et al.,1995). Ortiz-Lazareno et al. (2008) have demonstrated the effect of MG132 on the suppression of proinflammatory cytokine IL-6, TNF-_α_ and IL-1-beta release in the LPS stimulated human monocytic cell line U937.

## 5. Conclusion

Inhibition of proteasome function by PI plays an important role in the induction of apoptotic signals and anti-inflammatory responses in macrophages. The present study indicates that AM114 has a capacity to induce macrophage apoptosis and thereby can be considered as a therapeutic option. Altogether, our results showed that AM114 could induce apoptosis and decrease the release of proinflammatory cytokine and chemokine secreted by IL-1-beta stimulated THP-1macrophages. The results of this study suggest the use of AM114 as a therapeutic candidate in inflammatory diseases.

